# A three-dimensional musculoskeletal model of the dog

**DOI:** 10.1101/2020.07.16.205856

**Authors:** Heiko Stark, Martin S. Fischer, Alexander Hunt, Fletcher Young, Roger Quinn, Emanuel Andrada

## Abstract

Dogs are an interesting object of investigation because of the wide range of body size, body mass, and physique. In the last several years, the number of clinical and biomechanical studies on dog locomotion has increased. However, the relationship between body structure and joint load during locomotion, as well as between joint load and degenerative diseases of the locomotor system (e.g. dysplasia), are not sufficiently understood. In vivo measurements/records of joint forces and loads or deep/small muscles are complex, invasive, and sometimes ethically questionable. The use of detailed musculoskeletal models may help in filling that knowledge gap. We describe here the methods we used to create a detailed musculoskeletal model with 84 degrees of freedom and 134 muscles. Our model has three key-features: Three-dimensionality, scalability, and modularity. We tested the validity of the model by identifying forelimb muscle synergies of a beagle at walk. We used inverse dynamics and static optimization to estimate muscle activations based on experimental data. We identified three muscle synergy groups by using hierarchical clustering. Predicted activation patterns exhibited good agreement with experimental data for most of the forelimb muscles. We expect that our model will speed up the analysis of how body size, physique, agility, and disease influence joint neuronal control and loading in dog locomotion.

## Introduction

Dogs are an interesting object of investigation because of the wide range of body size, body mass, and physique of their more than 400 globally recognized breeds (Ostrander et al., 2019).

There exists an important body of work related to kinematical and dynamical differences between healthy dogs and dogs with musculoskeletal diseases (e.g. Brebner et al., 2006; Bockstahler et al., 2007; Burton et al., 2008; Burton et al., 2011). However, the relationship between body structure and joint load during locomotion, as well as between joint load and degenerative diseases of the locomotor system (e.g. dysplasia), are not sufficiently understood. We aim to investigate how body size, physique, agility, and diseases influence joint control and load in dogs.

To analyse joint mechanics, inverse dynamic analysis is typically used (e.g. Bresler & Frankel, 1950; Eng & Winter, 1995; Witte et al., 2002; Andrada et al. 2013). The inverse dynamics analysis is a method of the engineering sciences that combines kinetic, kinematic, and morphometric data to provide an indirect way to describe the causes of movement patterns. In order to quantify the joint load, the internal transmission of force through the skeleton, and consequently the generation of force in the muscles is required (Shahar et al., 2003). Simulated models, rather than invasive methods, are best suited to evaluate the force transmission between segmental elements (Shahar & Banks-Sills, 2002; Nielsen et al., 2003; Shahar et al., 2003; Shahar & Banks-Sills, 2004; Nyakatura & Andrada, 2013; Headrick et al., 2014).

Developing a model to test a scientific hypothesis leads to the question of its degree of complexity (Mehta et al. 2017). Interest in the general behaviour of the whole system (global dynamics) is a different approach than the analysis of joint mechanics or joint load. Thus, one could switch from simple models such as the spring-mass-model (Blickhan, 1989), to more complex multi-body models, or detailed models of body parts using the Finite Element Method (FEM). Model parameters (constant quantities during the simulation, e.g. mass or geometry), and model variables (speed, forces), must be obtained from experiments, literature, or ‘educated guesses’. Thus, the availability of model parameters and variables can influence the model’s complexity. In general, simple models (also termed templates by Full and Koditschek, 1999) are very well suited to study the basic principles of movement, while more complex models (termed anchors by Full and Koditschek, 1999) provide more realistic insights. A mixture of both extremes can be used to break-down the multidimensionality of complex models.

Two methods are distinguished to run simulations (Fig. 1). The forward simulation calculates specific torques in the joints on the basis of innervation data, EMG data, or muscle forces. Kinematics and finally locomotion (interaction between internal deformations with the environment) are derived. An inverse simulation, in which the joint torques are calculated from kinematic and kinetic data, determines muscle forces, EMG data, or innervation data a posteriori via static or dynamic optimization. The transformation of a one-dimensional variable (i.e. torque) into a multidimensional variable (i.e. muscle forces) is determined by numerical approximation. This means that for a given joint kinematics and joint torque, the muscle forces solution is not unique (underdetermined system).

**Figure 1:**
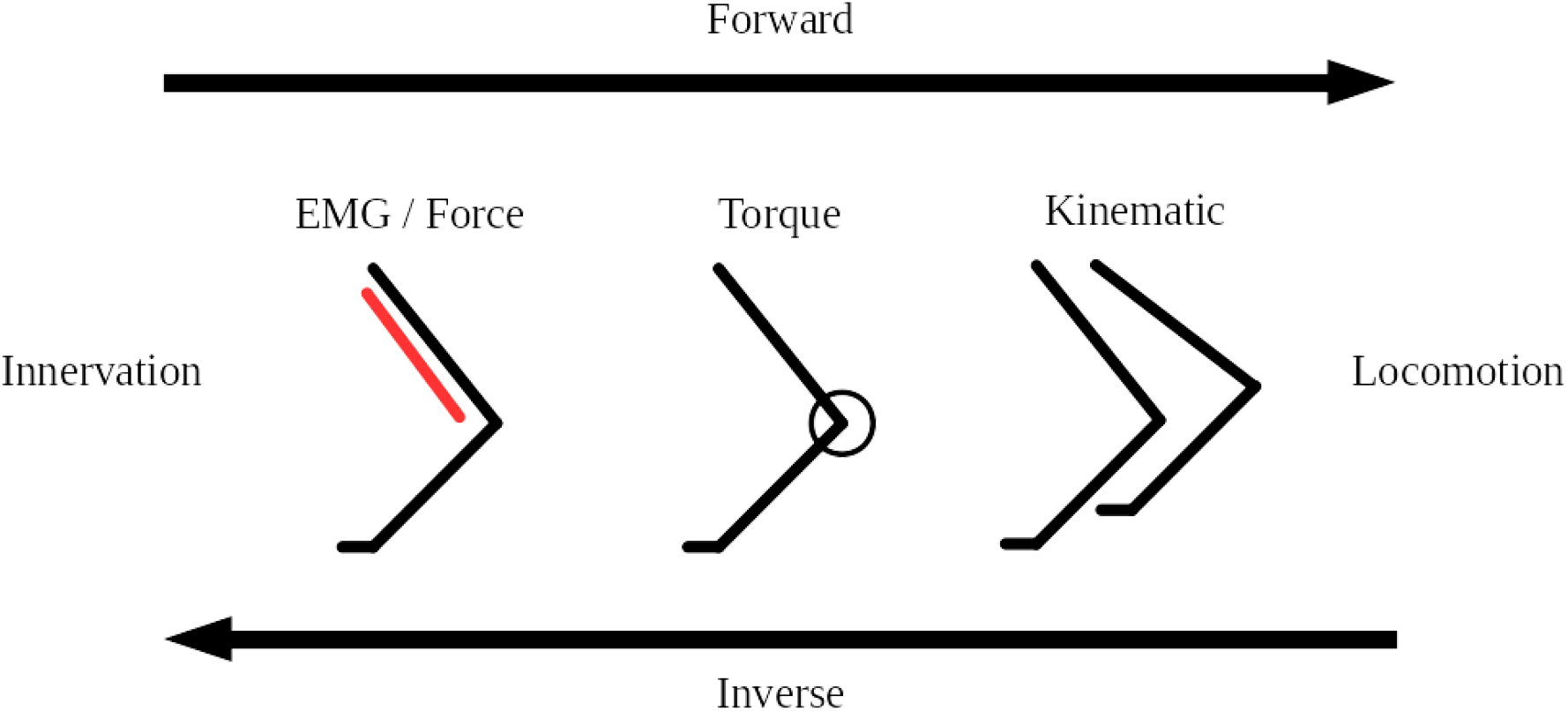
Abstraction of the forward and inverse simulation parameter chain for locomotion. Depending on the direction of the examination, the chain starts on the left or right side.

### Aim

We describe here the methods we used to create a detailed musculoskeletal model of a specific dog breed, the Beagle. Beagles are generalists that are often used for experimental purposes and in veterinary education. To permit a broad use of the model, we identified three key-features: Three-dimensionality, scalability, and modularity. Three-dimensionality is needed to represent a variety of movements such as periodic and non-periodic locomotion, agility, or idiomotion (e.g. scratching). Scalability is important because even within a dog breed, body, and limb proportions vary, but with some restrictions. Scalability also allows that the model should be useful to assess different breeds. Modularity helps to adapt the model to requirements/limitations of the experimental setup (e.g. single leg or multi-leg analysis).

A second, and specific goal of the present work is the identification of muscle synergies for a defined limb motion based on model simulations. This avoids invasive methods to record muscle activation and to include small and deep muscles into consideration. For this, we used inverse dynamics and static optimization to estimate muscle activations based on experimental data of a walking beagle. We present joint torques, muscle activations, and synergistic muscle groups for the forelimb. Torque profiles and muscle activation are then compared with published results.

## Material & Methods

### Computed tomography data

The Beagle model (BE-model) was built using computed tomography (CT) data of an adult Beagle (13.8 kg; Andrada et al., 2017). The resulting CT data set consisted of 3370 (spacing 0.33 mm) sections with a resolution of 512×512 pixels (spacing 0.279×0.279 mm). All skeletal bones were reconstructed from the CT data (Fig. 2). Care was taken to obtain an exact representation of the muscle attachment points on the bones. This simplified the precise transfer of postural and muscle data from another dog breed.

**Figure 2:**
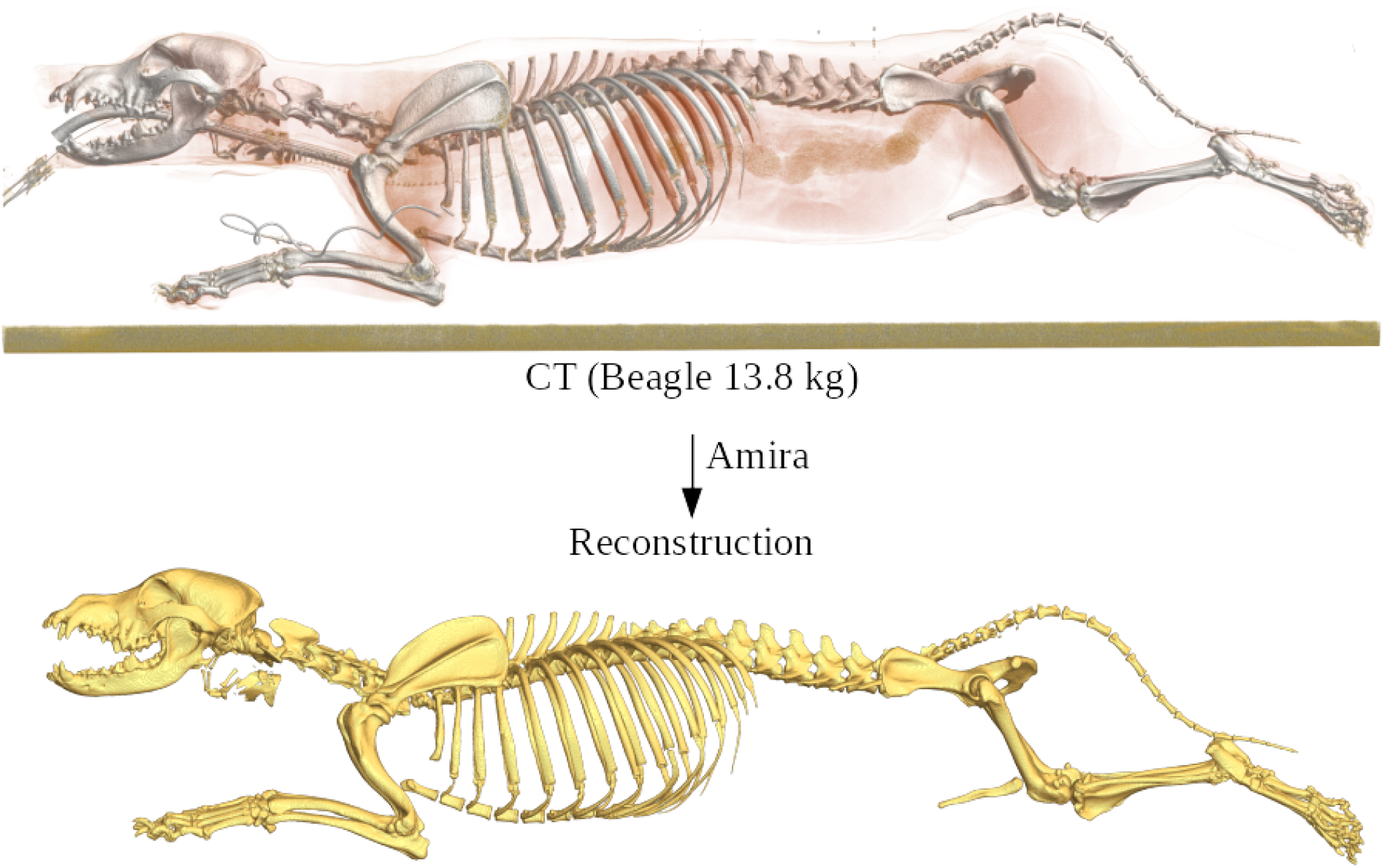
High-resolution CT data set of an anesthetized beagle (upper image) and the reconstruction of the separated bones using the software Amira® (lower image).

For reconstruction, we use the segmentation software Amira® [amira] and the analysis software imageXd [imageXd] for automatic mesh generation. In addition, the segment masses (Andrada et al., 2017) and their moments of inertia were determined from the CT data (see Suppl. Mass/Inertia).

### Muscle data

Because we do not have a musculoskeletal reconstruction of the beagle, we used an existing detailed anatomical model of the working line of the German Shepherd dog (GS-model) by J. Lauströer, A. Andikfar & M.S. Fischer (Fischer & Lilje, 2011) and transformed it to the beagle using the beagle’s skeletal model and insertion points. The GS model is based on cross-sections of the limbs and body as well as macroscopic dissections. The GS-model was created originally for illustration and animation in Autodesk Maya® [maya], and cannot be directly used in simulation tasks. Muscle paths were modeled as nurbs (non-uniform rational basis spline), which were spatially aligned transversely to the fibre course. The reconstruction of the muscle centrelines from the nurbs was performed in ‘Cloud2’ software [cloud2] (Fig. 3).

**Figure 3:**
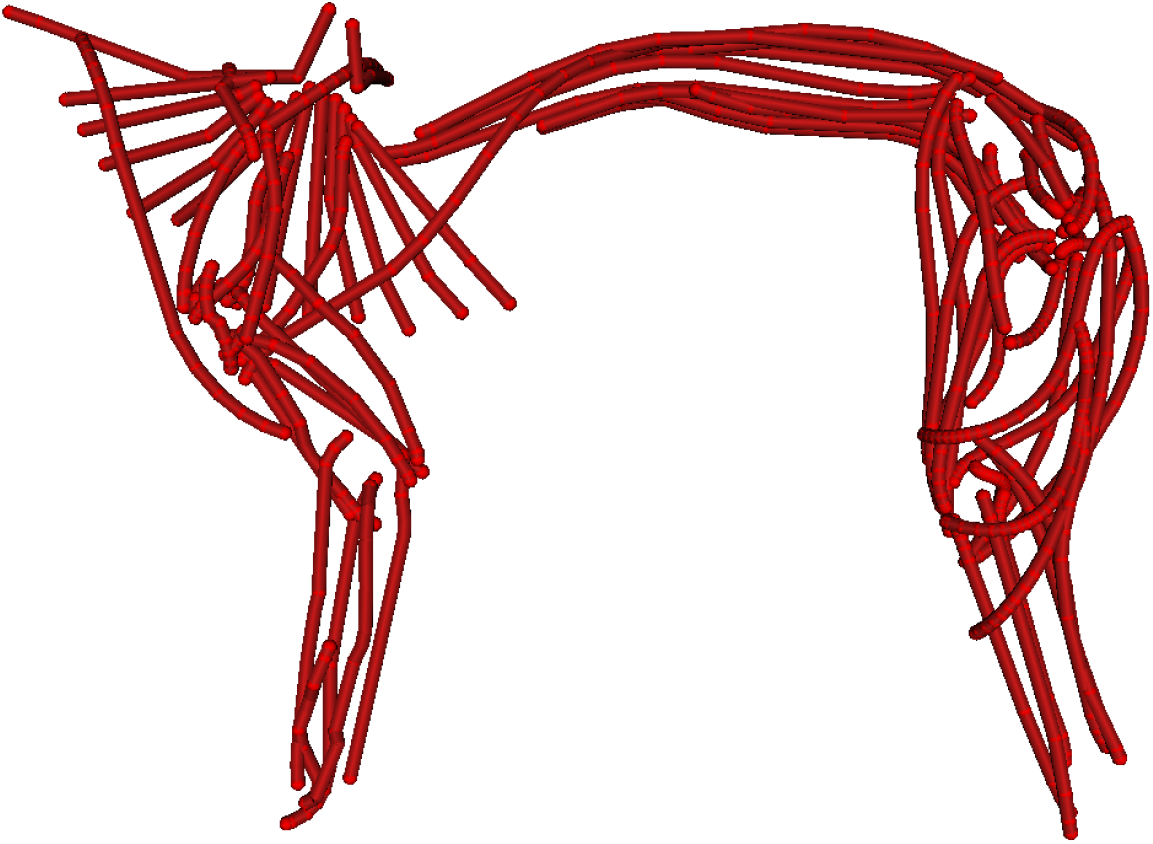
Muscle reconstruction based on the German Shepherd dog (GS) model for the fore and hind limbs as well as some trunk muscles.

The muscle parameters were taken from Shahar & Milgram (2001, 2005) and Williams et al. (2008a, 2008b). Shahar & Milgram published morphometric data of one hindlimb and four forelimbs of mixed-breed dogs. The morphometric variables included the muscle mass (*m*), muscle length (*ml*), muscle fibre length (*fl*), angle of pennation (*α*), and the resulting physiological cross-sectional area (*PCSA).* In addition, Williams et al. published morphometric data of seven forelimbs and six hindlimbs of racing Greyhounds. Here the morphometric variables included muscle mass (*m*), muscle length (*ml*), fascicle length (*fl*), as well as PCSA, maximum force, and power. The parameter of tendon length (*tl*), which is important for the model (Fig. 4), was not available and thus approximated using the following formula:

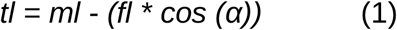

**Figure 4:**
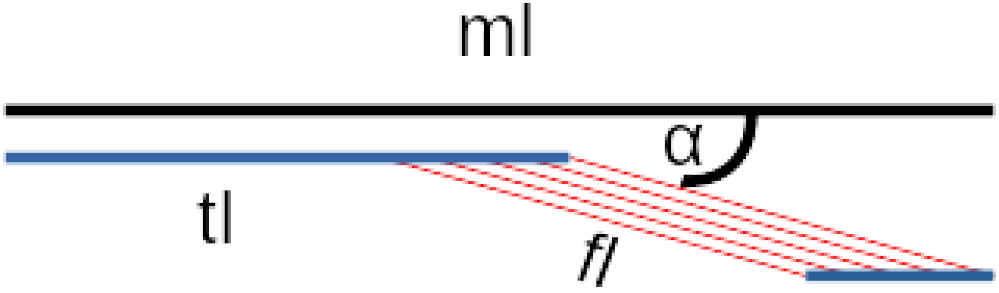
Illustration of the muscle parameter set for the tendon length calculation.

Adaptation or scaling of muscle parameters from other breeds or species is always a compromise. For the GE-model we used the muscle parameters from Shahar & Milgram (2001, 2005) and Williams et al. (2008a, 2008b). To scale those parameters for the beagle model (BE-model), we tested whether mass and total leg muscle PCSA scales with body mass for the published data. We found logarithmic relationships between both mass and total PCSA for a limb and body weight. Those relationships were used to scale every muscle PCSA to our BE-model (Fig. 5 & 6). Parameters that scale with length (e.g. muscle fiber, tendon slack length) are automatically scaled with the geometric change in OpenSim [opensim].

**Figure 5:**
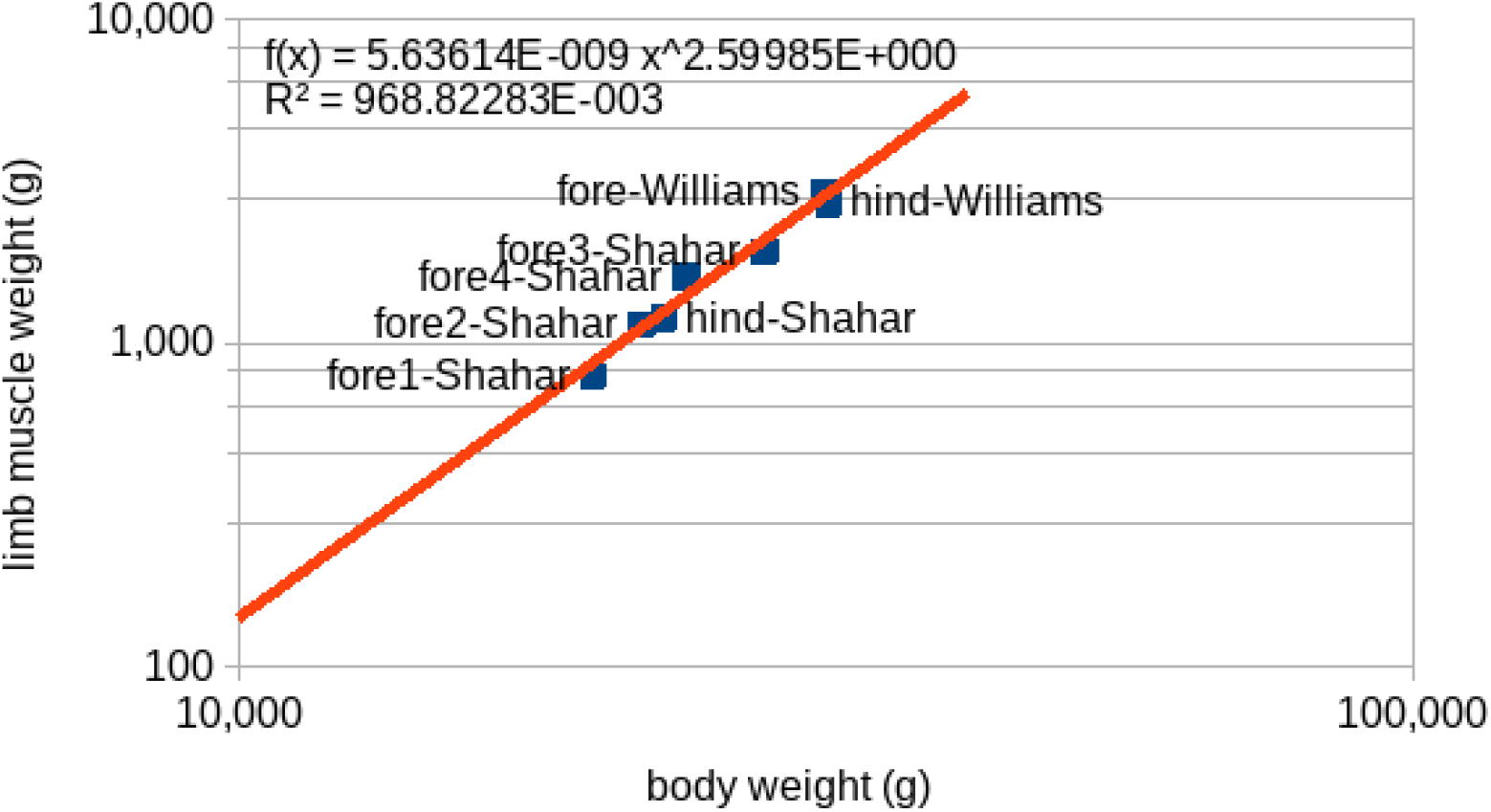
Comparison of the different total muscle masses and body weight between the dogs/studies (log. scale) for Shahar & Milgram (2001, 2005) and Williams et al. (2008a, 2008b).

**Figure 6:**
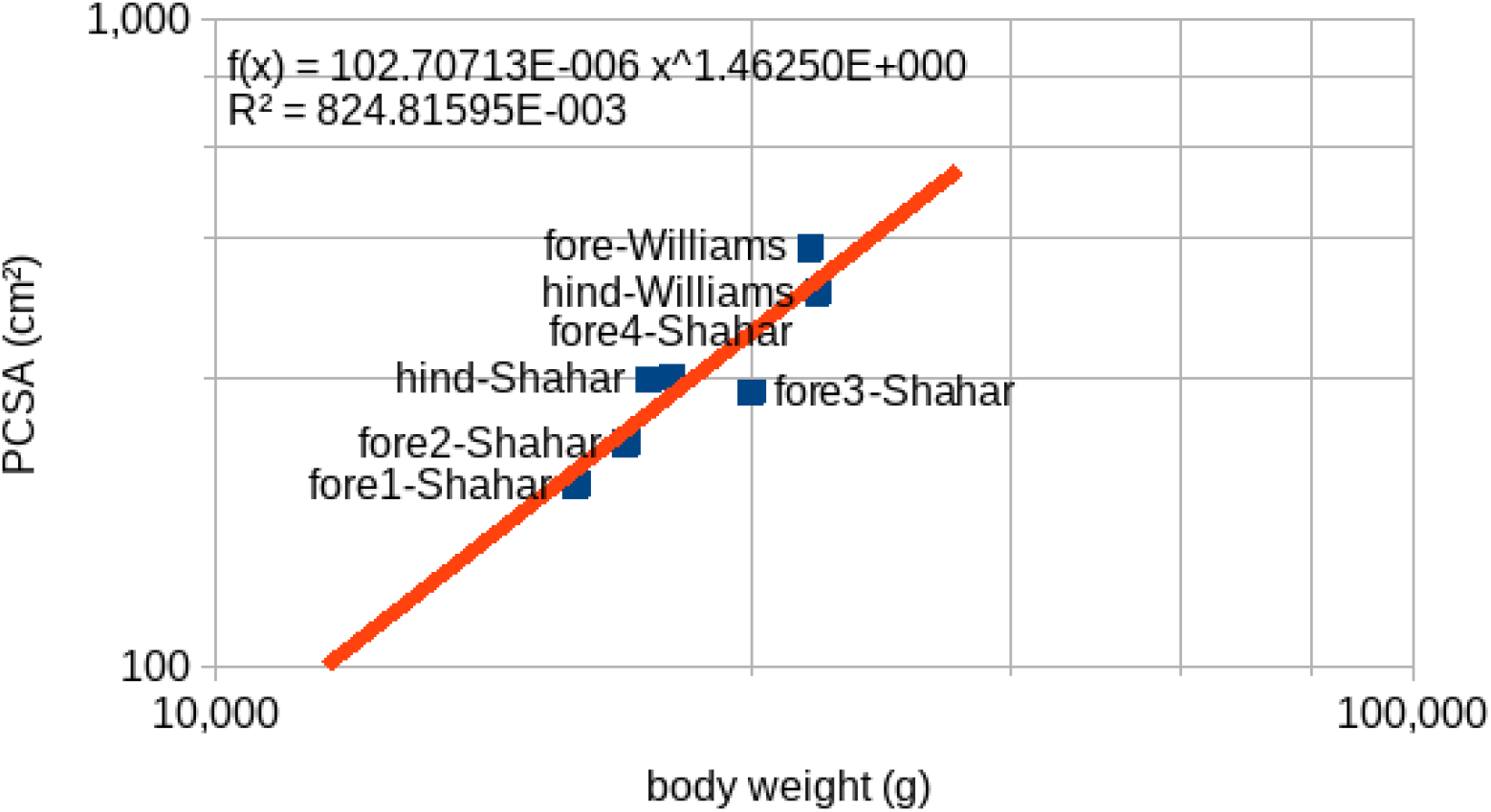
Comparison of the physiological cross-sectional areas (PCSA) and body weight between the dogs/studies (log. scale) for Shahar & Milgram (2001, 2005) and Williams et al. (2008a, 2008b). We used muscle parameters from different dog breeds. Because total leg muscle PCSA scales with body mass, we have used the same relationship to scale every muscle PCSA to our BE-model.

### Model assembly

To permit higher flexibility and broader use of the model, we generated the model in a way that it can be compiled in different languages. The basic script was written in Master [master], compiled as SIMM language [simm], and then converted via the simmToOpenSim tool [opensim] into the OpenSim language [opensim]. In addition, the scripts were created in such a way that we can flexibly create models with different specifications. Depending on the necessity, we can create the whole dog model or parts of it such as fore- or hindlimbs.

In the scripts, the segments are arranged hierarchically to build a kinematic chain as displayed in figure 7. The most proximal segment is joined to the ground (the thorax in case of the whole). The scripts include additional data for the relative position and orientation of the segments (bones), the segment masses, the centre of mass, and the inertia (bones-model). The individual sub-models (fore- or hindlimbs) contain only the mass of their corresponding segments.

**Figure 7:**
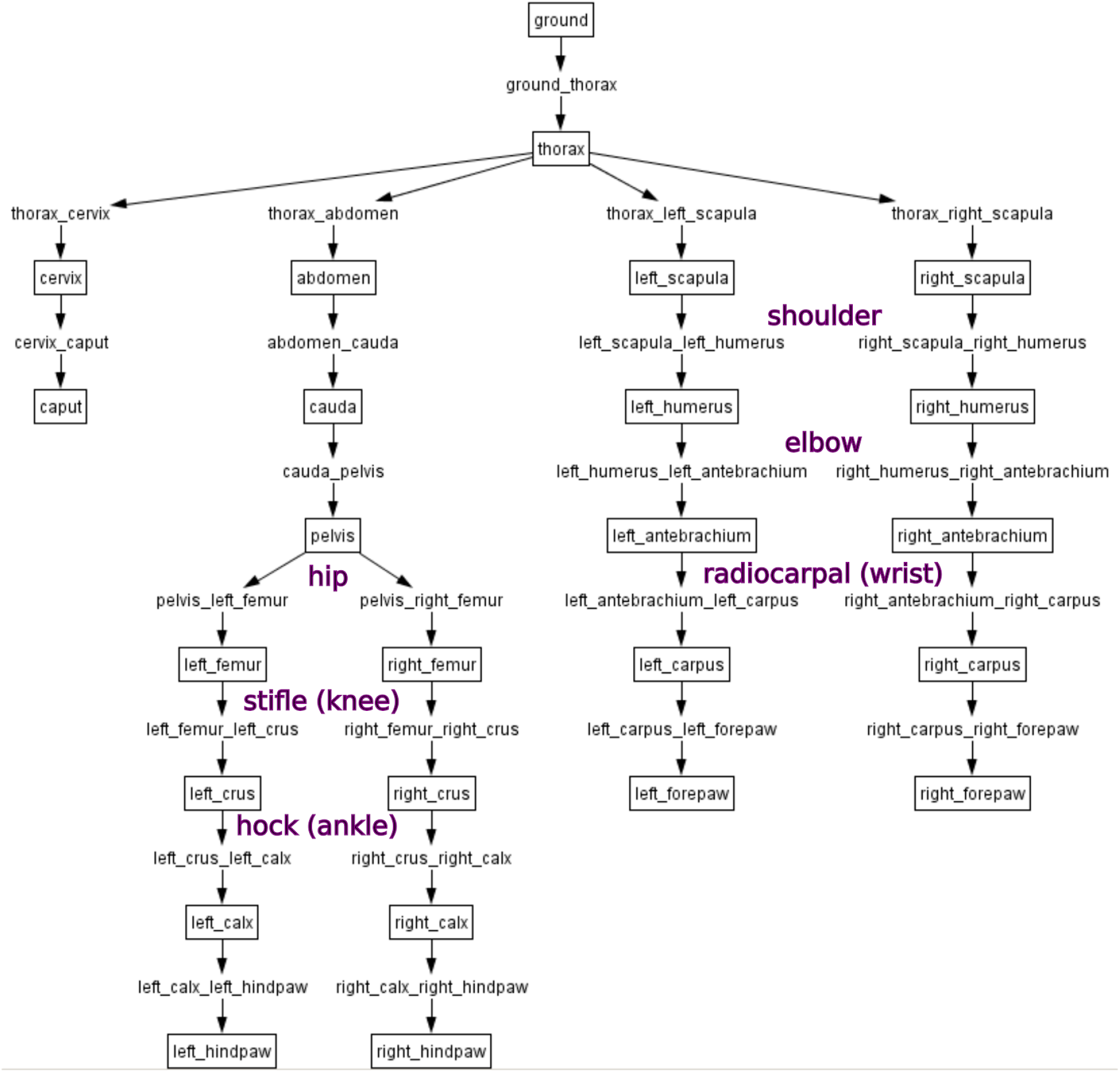
Topology of the segments (boxes) and joints (arrows) of the whole Beagle (BE) Model.

The bones-model (Fig. 8) is scaled and oriented to fit the muscle model (based on the GS-model). This task was performed in Blender ® 2.7. We scaled the bones model to fit the muscle model because this is easier to do than to scale muscles to fit the bones. In dogs, the segment length as a percentage of leg length is approximately the same among different dog breeds (Lumer, 1940; Wayne, 1986; Fischer & Lilje, 2011). Thus, just one size factor is necessary to scale a leg (in our case forelimbs and hindlimbs were multiplied by 1.66). By scaling beagle bones to fit the GS-model, muscle origins, and insertions of the muscle model (see Fig. 8) matched those of the Beagle bones’ model. Only small corrections on pelvis and scapula were necessary (see Suppl. Fig. S1). In the basic Master script, scaled BE-model and muscle-line models were combined. The position of the joints, position relative to the joint center, and joint types were derived from the CT-based BE-model. In addition, geometrical constraints have been added in the joints to prevent bone penetration by the muscles. Cylinders were used to constrain hinge joints while spheres were used to constrain ball-and-socket joints (Fig. 9). Our script permits us to generate either curved or simply linear muscle geometries. Muscle parts are assigned to their corresponding segments. Model segments (bones and muscle insertion points) can be then compiled in SIMM in global or local coordinates.

**Figure 8:**
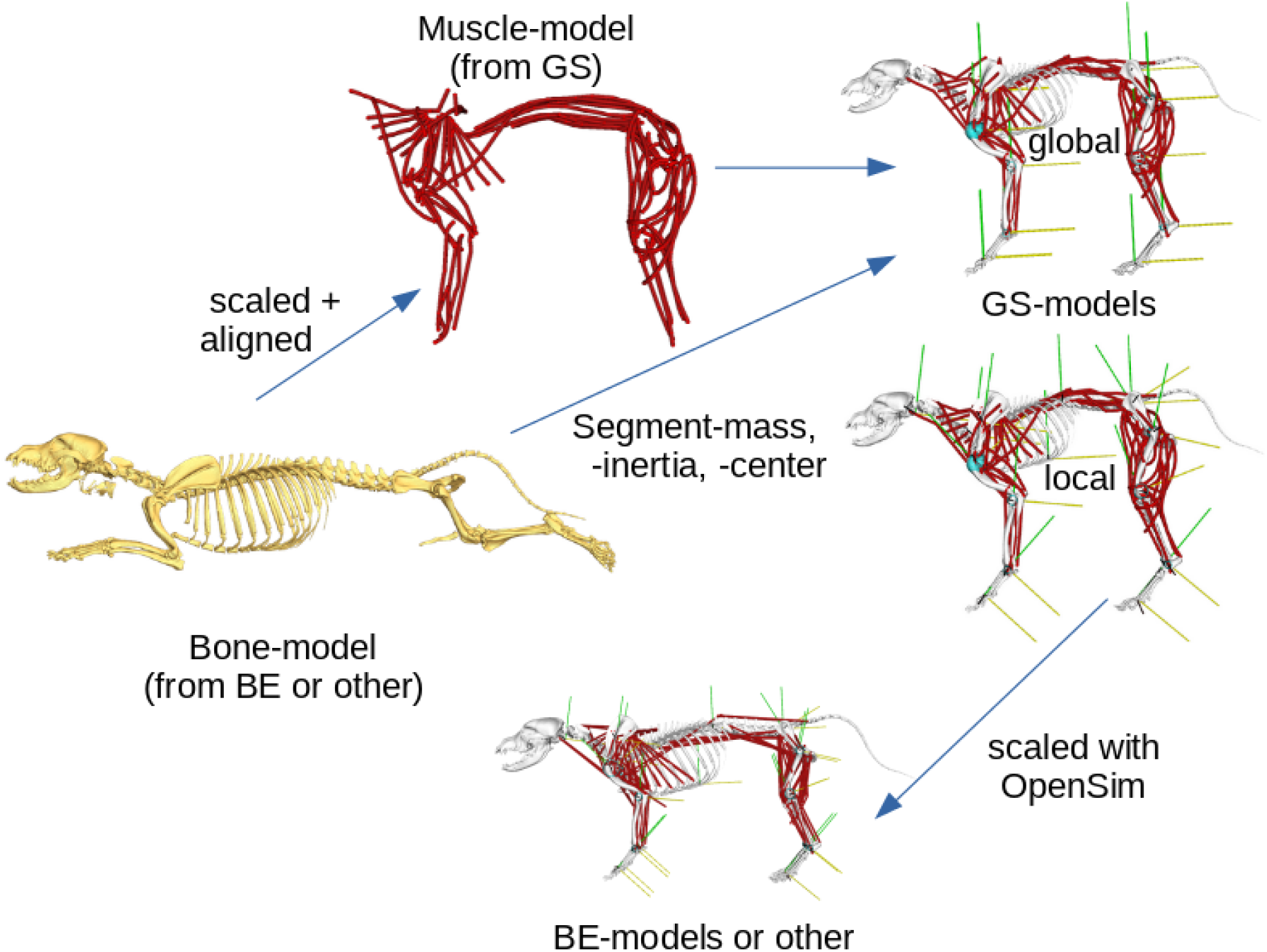
Representation of the model assembly, from the bone model to the muscle model to the resulting simulation model, taking into account the transformations performed.

**Figure 9:**
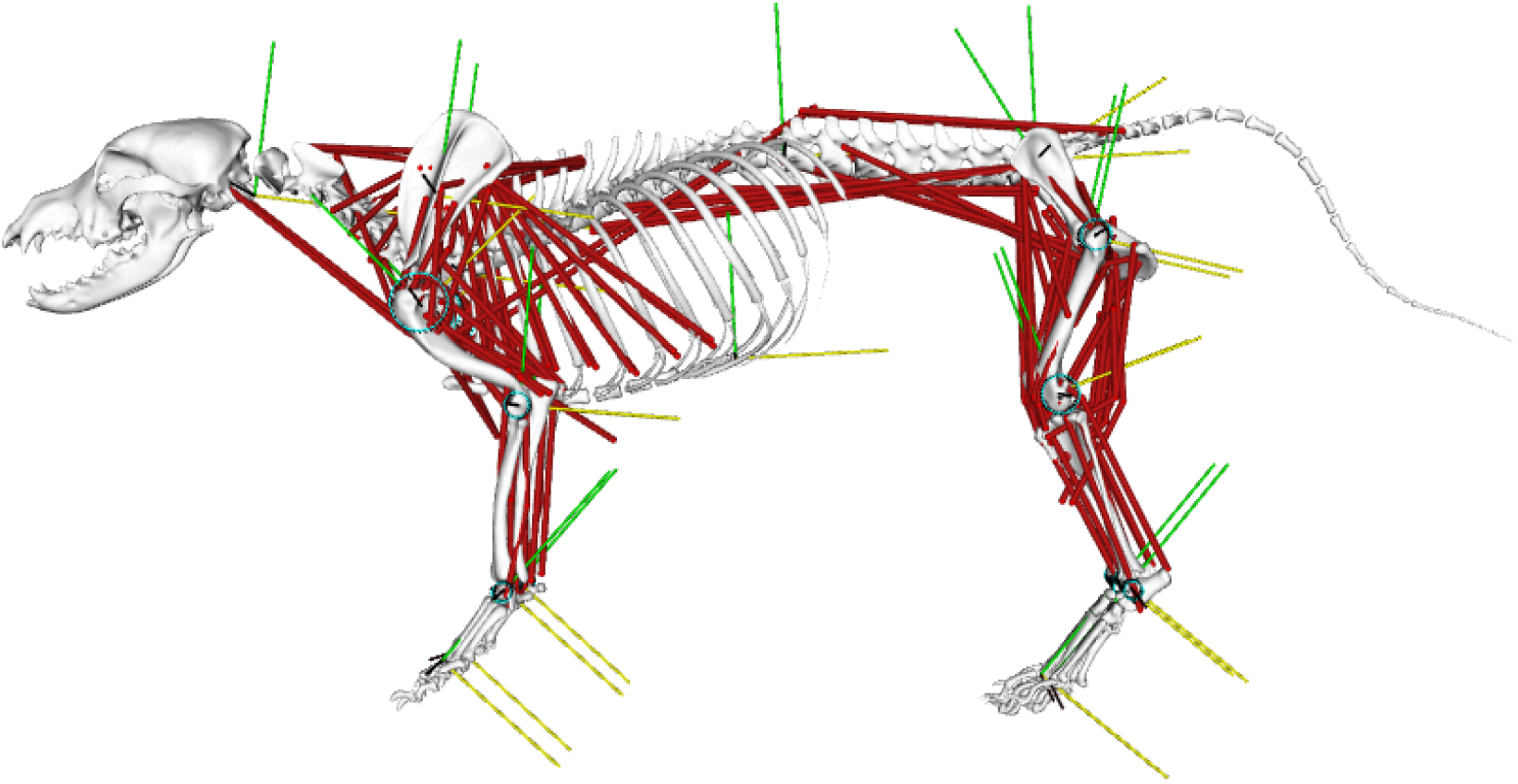
Full simulation model (BE-model with linear muscles) with visualization of the local coordinate systems (black, green, yellow) and the constraints (cyan).

### Experimental data, kinematics, GRF, and joint torques

In order to drive the simulations, we used 3D-kinematic and kinetic data from a previous dog study (Andrada et al., 2017). Ten walking strides (joint angles and ground reaction forces from left fore- and hindlimbs) from the same individual from which we took the CT scan were used as the basis for our analysis. We first computed 3D-Kinematic (XYZ Cardan-sequence) motion from marker data relative to the lab-frame. Because OpenSim necessitates 3D-relative joint coordinates, we transformed segment kinematics to joint kinematics by using quaternions. For this, we first transformed 3D-segmental Cardan angles to quaternions (for formulas see Henderson, 1977). Then, we computed the quaternions between adjacent segments. To do this, we conjugated the quaternion of the distal segment and then multiplied it by the quaternion from the most proximal segment. Afterward, we transformed the results again to relative Cardan angles XYZ.

For the sake of comparison, joint torques were also computed by using the 3D-Newton-Euler method (Eng and Winter, 1995; Winter, 2009), as already presented in Andrada et al., 2017. Here results are presented in the global frame to allow comparison with OpenSim (see supplementary document for more information).

### Joint actuators

We placed actuators in all joints to ensure that the muscle static optimization converges. This is important for two reasons. First, joint actuators ‘absorb’ numerical and mass errors. The second was to prevent those muscles from exceeding their maximal force. By scanning the parameter space the respective optimal joint actuator configuration was determined. Optimal parameters per joint were assumed when both the muscles spanning the joints stayed below their maximum force and the actuation was minimal. Actuator values can be found in table 1.

**Table 1:** The force and torque actuator values of the joints of the left forelimb calculated with a parameter map. As well as the degrees of freedom in the joints.

|  | **global** | **scapula** | **shoulder** | **elbow** | **carpal joint** | **forepaw** |
| --- | --- | --- | --- | --- | --- | --- |
| **$M_x$ [Nm]** | – | 5 | 0.0005 | 1 | 5 | 0.005 |
| **$M_y$ [Nm]** | – | 5 | 5 | 5 | – | 0.005 |
| **$M_z$ [Nm]** | – | 5 | 5 | – | 1 | 0.0005 |
| $F_x$ [N] | 0.5 | – | – | – | – | – |
| $F_y$ [N] | 0.5 | 0.00005 | – | – | – | – |
| $F_z$ [N] | 0.5 | 0.5 | – | – | – | – |
| **DOFs** | 3 | 5 | 3 | 2 | 2 | 3 |

### Simulation

To evaluate the model, we estimated inverse dynamics and muscle activation patterns for the forelimbs. Forelimbs are more challenging to model due to the high mobility of the scapular joint. Whereas the hindlimbs are linked to the pelvis via a locally static ball-and-socket joint, the scapula is not anchored to the body as a defined joint but via a complex arrangement of extrinsic appendicular muscles. Rather than rotating about a fixed point, scapular rotation incorporates both translation and rotation around an instantaneous centre of rotation.

We used OpenSim’s ‘inverse dynamics’ tool to compute the torques in the joints from the kinematic and ground reaction forces. With the ‘static optimization’ tool, we estimated muscle activation patterns and forces. We used the standard tool configuration (default) and the cut-off standard filter configuration (6 Hz) for the kinematic and ground reaction forces. We then compared the torque and muscle activation results with data from the literature.

Our goal was to reproduce muscular activation patterns of dog walking with the minimal possible DOFs ‘on’ in every forelimb joint. We started with a sagittal model (every joint represented as a hinge-joint). We then mapped different actuator force combinations ranging from 1E-9 to 1E9. We analyzed the simulated muscle activations and compared them to the results in the literature. We then increased one DOF in one joint on one plane from the most distal to the most proximal one and mapped again the optimal set of muscle actuators. This procedure was repeated in every joint until the increment of a joint-DOF did not improve simulation results.

### Hierarchical cluster-analysis

In order to evaluate synergistic muscle groups, the predicted muscle activations were further analyzed using the hierarchical cluster analysis. This method is typically used to find and group similar patterns within a data set. We first determined the Euclidean distance between the time-series datasets of logarithmic muscle activations. The Euclidean distance matrix over time was then used to group the data and generate a similarity tree, which was again sorted based on the minimum distance. To perform this analysis we used the software package R [r] (packages: dendextend, ggdendro and dendsort).

## Results

### Dog model

The complete model has a maximum of 84 DOF. Head, thorax, abdomen, and tail have three rotational DOF each. Vertebral motions were not modeled. We included 134 (67 per side) muscles (with their corresponding muscle parameters) encompassing the majority of fore- and hindlimb muscles and the epaxial muscles relevant for locomotion. Head, belly, and toe muscles were not included. The complete set of muscles can be found in (Suppl. Tab. S1).

### Left forelimb

To evaluate the model, the left forelimb was chosen as the test case. For simplicity, linear muscles were used. Note that ‘linear’ does not include the final end of the muscles. There, if necessary, muscles followed the geometric constraints modeled to avoid bone penetration. As expected, the number of degrees of freedom per joint influenced the results of the muscle activation. For the forelimbs, 15 DOFs were necessary to converge static optimization to muscle published activations: scapula (5 DOFs), shoulder (3 DOFs), elbow (2 DOFs), carpal joint (2 DOFs), paw (3 DOFs) (details see Tab. 1). For example, without scapular anterior-posterior translation, the activation of the *M. rhomboideus* and *M. latissimus dorsi* peak during the early stance phase. Such an activation profile has not been reported in the literature. Once the translational DOFs in the scapular joint were allowed, the *M. rhomboideus* was activated continuously, and the *M. latissimus dorsi* was active at the end of the swing phase in accordance with experimental observations (Tokuriki, 1973; Goslow et al. 1981; Deban et al., 2012). Increasing the DOFs to more than 15 did not significantly improve the simulation results.

### Torques

The comparison between torque results computed by OpenSim and those computed using the Newton-Euler method (see supplement for method details) show more agreement on the sagittal plane (Fig. 10). The frontal plane displays higher discrepancies (Fig. 10). In the horizontal plane, differences can be observed for the scapula and the humerus (Fig. 10). Joints showing larger differences in those two planes, display also the larger standard deviations in the result computed using OpenSim. Still, the torque amplitudes and patterns computed using OpenSim are similar to those computed using the Newton-Euler method and to those published (Nielsen et al., 2003; Andrada et al., 2017, Fig. 10). The thorax shows the largest torque amplitude in the sagittal plane, followed by the scapula and the humerus, which display similar torque patterns and amplitudes. In the frontal plane, besides the carpal joint and the forepaw, all torque patterns and amplitudes were similar. In the horizontal plane, the shoulder joint displays larger torque amplitudes.

**Figure 10:**
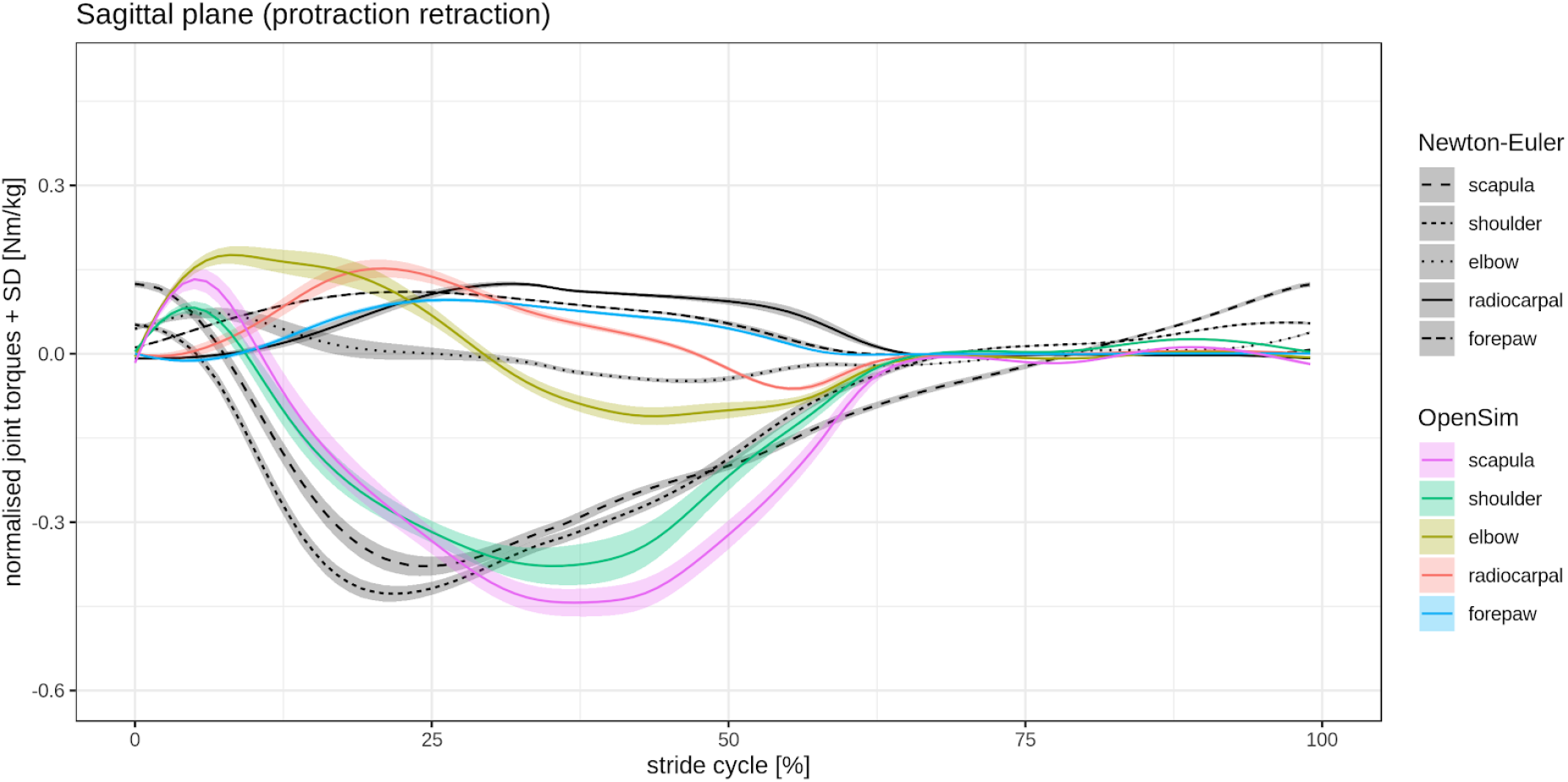
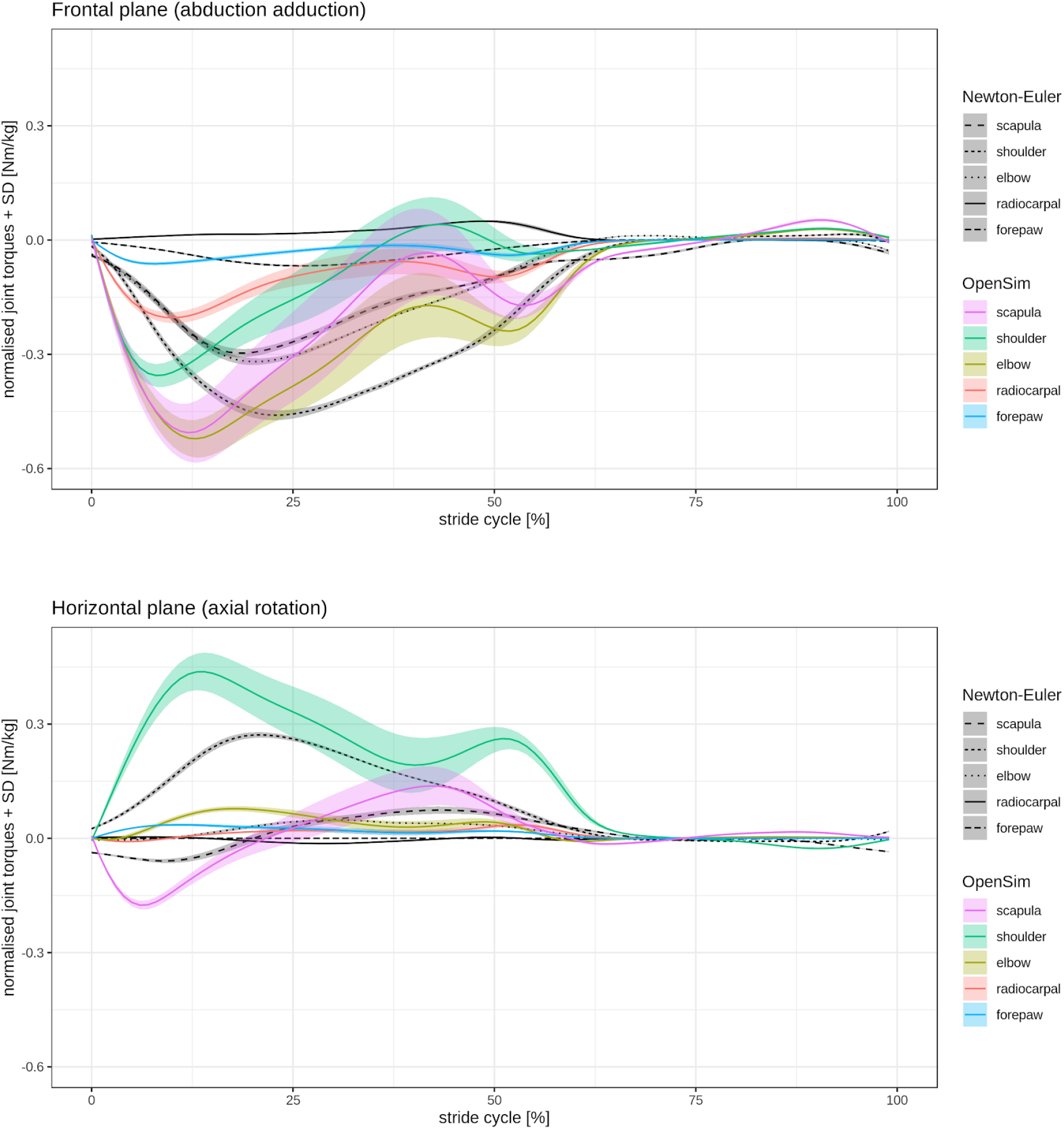
Forelimb calculated torques for the sagittal, frontal, and horizontal plane in a walking beagle. Comparison between results obtained from OpenSim vs. Newton-Euler method. The standard deviation (SD) is shown as shaded bands.

### Muscle activation

The results of the static optimization converged to muscle activations, which were similar to those collected in experiments (Tokuriki, 1973; Deban et al., 2012). The muscles that exhibited the largest activations were the *M. supraspinatus* and the *M. infraspinatus* (Fig. 11; Suppl. Fig. S2). Both muscles were activated during the whole stance phase. Interestingly, all other muscles were activated either during the early or late stance phase. For muscles in this stance boundary group, the *M. serratus* displayed an activation wave traveling from their cranial to their caudal parts. Note that the cranial parts retract, and the caudal parts protract the leg.

**Figure 11:**
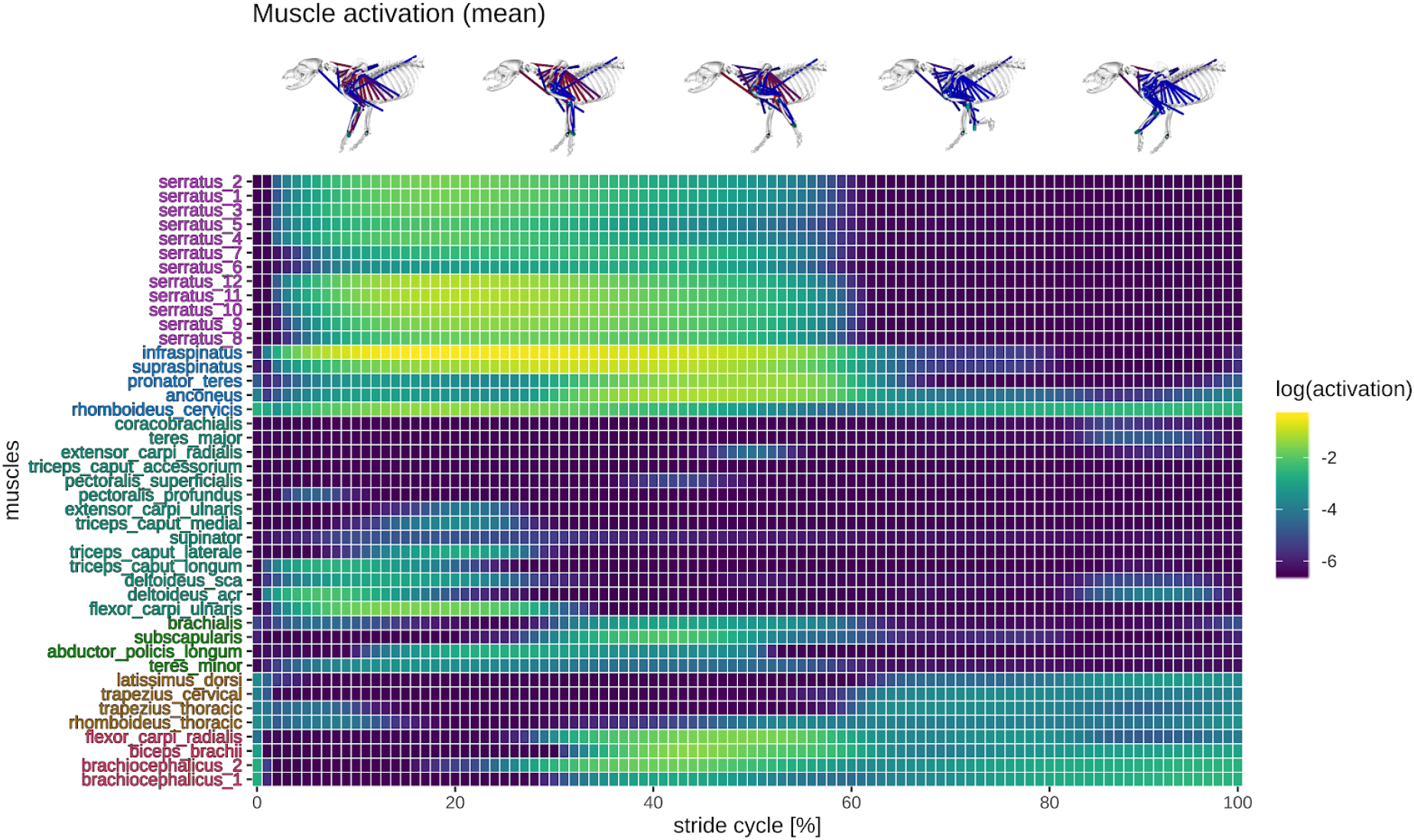
Simulated forelimb muscle activation in a walking beagle. Colors represent muscle activation levels on a logarithmic scale.

The hierarchical cluster-analysis separated muscles into three main synergistic groups (Fig. 12 – black lines; Suppl. Fig. S4). Every main synergistic group has two subgroups (see different colors in Fig. 12). The purple subgroup of group #1 includes all of the different parts of *M. serratus.* To the subgroup blue of group #1 belong *M. infraspinatus, M. supraspinatus, M. pronator teres, M. anconeus, and M. rhomboideus cervicis*.

**Figure 12:**
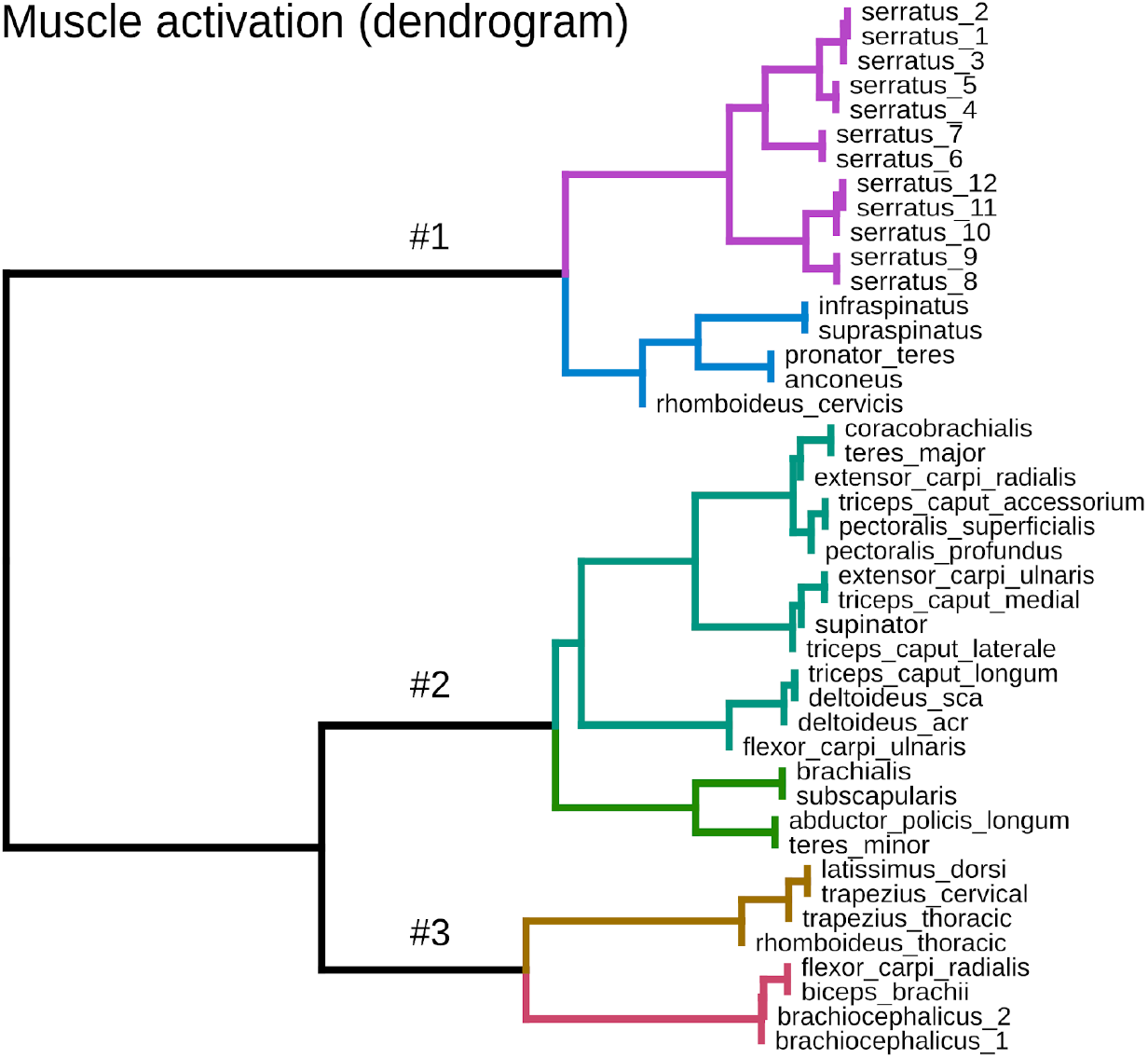
Hierarchical clustering (method – ward.d2) and minimal leaf sorting of simulated forelimb logarithmic muscle activation in the walking beagle.

The cyan subgroup of group #2 encompasses a large number of joint extensor and joint stabilizing muscles. The first group encompasses the *M. coracobrachialis*, both parts of *M. pectoralis*, *M. triceps accessorium, M. teres major*, and *M. extensor carpi radialis*. The second group of muscles within the cyan subgroup include: *M. extensor carpi ulnaris, M. supinator, M. triceps caput longum, caput medialis, and caput lateralis, M. deltoideus*, and *M. flexor carpi ulnaris.*

The brown subgroup of group #3 include the *M. latissimus dorsi, M. trapezius pars cervicalis*, *M. trapezius pars thoracis*, *M. rhomboideus pars thoracic*. The magenta subgroup of group #3 contains the *M. flexor carpi radialis*, *M. biceps brachii*, and both parts of the *M. brachiocephalicus.*

## Discussion

Dogs have more than 400 globally recognized breeds (Ostrander et al., 2019). Thus, they permit a unique possibility to analyse how body size, physique, and agility, as well as diseases, influence joint control in quadrupedal locomotion. The aim of this work was to develop a detailed, fully three-dimensional, and scalable musculoskeletal model of a dog to analyze these effects.

We designed a flexible framework to generate a whole or parts of dog musculoskeletal models of different breeds. To our knowledge, our model is the first complete and fully three-dimensional model of a dog having 134 (67 per side) muscles including the majority of fore- and hindlimb muscles and those epaxial muscles relevant for locomotion.

To evaluate the model, we estimated inverse dynamics and muscle activation patterns for the left forelimb. We compared the results of the inverse dynamic tool in OpenSim to the inverse dynamic computations based on the same data using the Newton-Euler method and published data.

### Joint torques

There exist only a few available studies on 2D-inverse dynamic analysis of the canine forelimb (i.e. excluding shoulder joint and scapular fulcrum, Nielsen et al., 2003; Burton, 2008; Burton 2011) and only one in three-dimensional analysis of the whole forelimb (Andrada et al., 2017). Note that only Niesen et al. (2003) and Andrada et al. (2017) reported on healthy dogs, and therefore the results of Burton, et al. (2008) and Burton et al. (2011) will not be further discussed here.

As a standard configuration, OpenSim presents joint-torques projected on global planes. For comparison, we projected joint-torques computed with the Newton-Euler method transformed to those same global planes. One can discuss whether this description is more or less useful. However, for the purposes of this work, we were interested in checking the functionality of the dog model. In general, the torque amplitudes and patterns computed using OpenSim were similar only in the sagittal plane to those computed using the Newton-Euler method and to those published (Nielsen et al., 2003; Andrada et al., 2017, Fig. 10). Differences in torque amplitudes were observed for the frontal and horizontal planes. Discrepancies between OpenSim and Newton-Euler methods are to be expected because of the error accumulation in the recursive Newton-Euler method and differences in filtering. We can not be sure how OpenSim understood the Cardan-sequence of the relative joint angles. Even small differences in leg orientation related to the force vector might also help to explain the discrepancies observed, especially in the frontal plane.

### Muscle activation patterns

OpenSim uses a two-step process to estimate muscle activation: First, inverse dynamics are used to compute joint forces and then static optimization is used to compute muscle forces. It has been argued that this methodology hampers OpenSim’s ability to correctly solve closed-loop systems (Cadova, 2013). Other software, such as AnyBody, can more accurately solve closed-loop systems because they solve for the inverse dynamics together with the optimization (i.e. muscle and joint forces are unknown at the same time). Introducing experimentally derived ground reaction forces, the OpenSim model is basically an open loop. Thus, the differences between AnyBody and OpenSim will lessen (Kim et al., 2018; Trinler et al., 2019).

For the simulations, we choose the Hill-type muscle model by Millard et al. (2013). We took muscle parameters from the works by Shahar & Milgram (2001, 2005) and Williams et al. (2008a, b) and linearly scaled them to the body mass of a beagle. Additionally, we estimated the tendon length (see methods). As a general rule, muscle contractions in OpenSim are not effective in stabilizing joints while also reproducing limb kinematics during walking. Therefore, we included actuators in all joints to assist the muscles. A parameter search found the minimal actuator force/torque values that permitted the static optimization to converge.

Model predictions of muscle activations, including the correct separation of the muscle groups with regard to the stance and swing phase, showed good agreement with reported EMG-data (Tokuriki, 1973; Goslow et al. 1981; Deban et al., 2012; Suppl. Fig. S5 & S6). Exceptions were the *M. triceps* and *M. trapezius* (compared to Tokuriki), the *M. pectoralis* (compared to Tokuriki, 1973 & Deban et al. 2012), and *M. serratus ventralis cervicalis* (compared to Deban et al., 2012). Muscle activation is sensitive to the assigned muscle characteristics and joint actuator parameters. A sensitivity analysis could provide insight into what specific muscle/joint properties have the most effect on muscle activation patterns. However, such an analysis is outside the scope of the present work.

Muscle activation patterns in the literature are sometimes inconsistent (*M. pectoralis profundus* => Tokuriki ≠ Deban; *M. latissimus dorsi* => Deban ≠ Tokuriki), making the comparison to a ground truth difficult. The position of the electrodes (especially in large muscles), time-varying activation of different muscle regions, and muscle cross-talk may explain these differences. This assumption can however not be tested, as e.g. Tokuriki (1973) did not report the position of the electrodes.

One important finding of this work is the relationship between joint DOFs and muscle activation. For example, predictions of the activation patterns of the *M. rhomboideus* and *M. latissimus dorsi* are similar to those reported in the literature only when the scapular joint has five or six DOFs. Anterior-posterior translation in the scapular joint must be present as observed in kinematic studies (see Fujiwara, 2018 for review). The fact that the scapula is only linked to the body with muscles indicates that the scapular joint might hold an additional role as a damper, reducing the propagation of impact forces to the body (e.g. after jumping) and minimizing the necessity of gait compensation mechanisms.

### Validity and muscles synergies

We used a hierarchical cluster-analysis to further analyze the validity of our musculoskeletal model. This analysis identifies synergistic groups by separating muscles with similar activation patterns. This method is widely used in the literature to classify motor neuron activation, movement, or disease-related differences (e.g. Pennartz et al. 1998; Mentel et al. 2006; Ferrarin et al., 2012).

The theory of the use of muscle synergies (e.g. D’Avella et al., 2003; Ting et al., 2007; Markin et al., 2012) hypothesizes that the CNS produces different motor behaviors by co-activating groups of muscles in space or time (D’Avella et al. 2003, Hart & Giszter, 2013). Two types of muscle synergies have been identified (Tresch & Jarc, 2009; Hart & Giszter, 2013): (1) ‘synchronous synergies’, which activate a group of muscles at the same time; and (2) ‘time-varying synergies’, which produce patterns with a temporal profile for each muscle of a synergistic group. Testing muscle synergies is a method to understand how the central nervous system (CNS) produces a wide range of motor behaviors and could be an important tool to simplify the control problem in complex neuromechanical models (Markin et al., 2016; Deng et al., 2019).

Synergy decomposition yielded three main groups. Every group was further divided into two subgroups, which have different colors (group #1: purple and blue, group #2: cyan and green, group #3: brown and magenta). The branch out among one color indicates some delay in the activation. The purple subgroup of group #1 belongs to all the different parts of the *M. serratus*. Those parts were activated sequentially from the most cranial to the most caudal parts. The cranial parts retract, while the most caudal parts protract the scapula. The protraction of the scapula correlates with the braking ground reaction forces observed in most of the stance phase, while the retraction correlates with the acceleration phase in the late stance.

The blue subgroup of group #1 includes *M. infraspinatus, M. supraspinatus, M. pronator teres, M. anconeus, and M. rhomboideus cervicis. M. supraspinatus* and *M. infraspinatus* extend the shoulder joint, *M. anconeus* extend the elbow joint, while *M. rhomboideus cervicis* mainly stabilize the scapular joint. In our simulations, they were recruited during stance and part of the swing phases. However, only the *M. rhomboideus cervicis* was active throughout the entire stride. EMG-data exists for the *M. supraspinatus*, *M. infraspinatus*, and *M. rhomboideus cervicis*. Our simulation results display a good agreement with those experimental data (Fischer & Lilje 2011, Tokuriki, 1973 & Deban et al. 2012).

The cyan subgroup of group #2 encompasses a large number of muscles. Among this subgroup, two main activation patterns were predicted. The first group encompasses the *M. coracobrachialis*, both parts of *M. pectoralis*, *M. triceps accessorium, M. teres major*, and *M. extensor carpi radialis*. Those muscles showed minimal activations during simulations, which differs from published EMG-data. One explanation for these differences is that more accurate muscle parameters or/and new goal functions for optimization are needed to better distribute force among muscles. A further explanation could be, based on the fact that these muscles are difficult to be measured, that the published data displayed just a cross-talk to more superficial muscles. Maybe these muscles are recruited for other tasks (e.g. perturbed locomotion).

The second group of muscles among the cyan subgroup (*M. extensor carpi ulnaris, M. supinator, M. triceps caput longum, caput medialis, and caput lateralis, M. deltoideus*, and *M. flexor carpi ulnaris*) were recruited in the early stance phase. They work mainly against gravity and control the axial function of the leg. The axial function refers to the time-dependent length and force of the effective leg, which is a model leg from the main proximal pivot/fulcrum (here scapular joint) to the foot (e.g. Maus et al., 2010; Andrada et al., 2014; Nanua & Waldron, 1995).

The green subgroup of group #2 is made up of the following muscles: *M. brachialis, M. subscapularis, M. abductor pollicis longus, M. teres minor.* In the literature there exists EMG-data only for the *M. brachialis*. Experimental data show that during walking, this elbow flexor is recruited from late stance until mid swing approximately (Fischer & Lilje, 2011). In our model, the *M. brachialis*, while displaying similar whole activation time, started about 10% of the stride time earlier than in the experiments. Interestingly, in our simulations, the *M. subscapularis* had a similar activation pattern to those predicted for the *M. brachialis.* The *M. abductor pollicis longus* and the *M. teres minor* were activated earlier in the stance phase. The former was activated around midstance, while the latter was activated during early stance phase. The *M. teres minor* displayed a similar activation pattern to the EMG-data published for the *M. teres major* (Fischer & Lilje, 2011), This could indicate that the *M. teres minor* took the place of the *M. teres major* in our simulations. However, the turning-off the *M. teres minor* in simulations did not significantly improve the predictions of the activations of the *M. teres major*.

The third group encompasses muscles that were activated throughout the swing phase. Some of them were also recruited during parts of the stance phase. The brown subgroup of group #3 includes the *M. latissimus dorsi) M. trapezius pars cervicalis, M. trapezius pars thoracis, M. rhomboideus pars thoracico* The predicted activation of *M. latissimus dorsi* resembles experimental findings. It brakes the protraction of the forelimb during swing before touchdown. On the other hand, the *M. trapezius* and *M. rhomboideus* stabilize the scapular joint. In experiments, the *M. trapezius pars cervicalis* was active during the entire stride cycle, while the *M. trapezius pars thoracis* was active during the complete stance phase and at the late swing phase (Fischer & Lilje, 2011). Our simulations predicted for the *M. trapezius* to be active earlier during stance and during the complete swing phases. EMG data shows that the *M. rhomboideus pars thoracis* is active during most of the stride cycle with the exception of a short period around midswing. In our simulations, this silent period occurs around midstance.

The magenta subgroup of group #3 contains the *M. flexor carpi radialis. M. biceps brachii*, and both parts of the *M. brachiocephalicus*. The first two muscles flex the paw and elbow joint, respectively, while the third protracts the forelimb. In the literature, it was shown that these muscles have similar activation patterns. They are briefly active after touchdown, then around take-off, and in the late swing phase. Our simulations display a similar pattern to those found in experiments. Furthermore, they show that the largest activations occur around take-off in preparation and start of the swing phase.

### Conclusions

We have developed a musculoskeletal model of a dog that has three main features: three-dimensionality, scalability, and modularity. Activation patterns predicted by static optimization exhibited good agreement with experimental data for most of the forelimb muscles. However, because muscles were unable to stabilize joints, joint actuators have been included. In animals, joints are stabilized by muscle co-contraction and passive structures (Dhaher et al., 2010; John et al., 2013; Flaxman et al., 2012; Knarr et al., 2012). Thus, muscle geometry, muscle parameters, and the modeling of passive structures are essential for an accurate estimation of muscle activation. To this end, more detailed breed-related anatomical and physiological studies are necessary. Other optimization algorithms as the default algorithm in OpenSim or predictive forward simulations might also improve the predictive power of this dog model (e.g. Geijtenbeek, 2019).

We expect that the use of our model will speed up the analysis of how body size, physique, and agility (as well as disease) influences joint control and loading in dog locomotion. We follow two different paths for the expansion of this model: a) we are modeling specific joints in more detail using the finite element method to analyse joint loads based on the force data of the current model; b) we intend to expand the model to a neuromechanical model (c.p. Hunt et al., 2015; Deng et al., 2019), to understand how the neural, muscular and skeletal systems operate together to produce efficient and stable locomotion. For this, we also present in this work a method to estimate muscle synergies, which can help to break-down the design complexity of neuronal networks.

## Acknowledgments

We thank all the involved members of the Institute of Zoology and Evolutionary Research for their stimulating discussions and support. We especially thank Jonas Lauströer and Amir Andikfar for providing the muscle geometry for the model. This research work was supported by the company ‘Biologische Heilmittel Heel GmbH’ and by grants from the US-German CRCNS program including DLR grant 01GQ1605 and NSF grant 1608111.

**imageXd** – (http://starkrats.de)

imageXd is a command-line tool to process and evaluate spatial data (such as CT or MRI data). Additionally, it can process vector-, tensor fields and mesh data.

*platforms:* windows, mac os & linux (free research software)

*version*: 4.6.13

**cloud2** – (http://starkrats.de)

cloud2 has a graphical user interface (GUI) and can process and evaluate data such as point clouds, vector fields, and space curves.

*platforms* : windows, mac os & linux (free research software)

*version* : 12.6.9

**Amira** – (https://www.fei.com/software/amira-3d-for-life-sciences/)

A commercial, high-level language and interactive environment for numerical computation, visualization, and programming.

*platforms*: windows, mac os & linux

*version* : 5.4

**Maya** – (https://www.autodesk.de/products/maya/overview)

Maya®, is software for animation, modeling, simulation and rendering in 3D, offering integrated powerful tools for animation, environments, motion graphics, virtual reality, and character creation.

*platforms*: windows, mac os & linux

*version*: 2016

**OpenSim** – (http://opensim.stanford.edu/)

OpenSim is a freely available software system that allows you to build, exchange, and analyse musculoskeletal models and dynamic simulations of movement.

*platforms*: windows

*version* : 3.3

**R** – (https://www.r-project.org/)

R is a free software environment for statistical computing and graphics.

*platforms*: windows, mac os & linux

*version*: 3.5.3

**SIMM** – (http://ftp.motionanalysis.com/html/movement/simm.html)

SIMM (Software for Interactive Musculoskeletal Modelling) is a powerful tool kit that facilitates the modeling, animation, and analysis of 3D musculoskeletal systems.

*platforms*: windows

