## Supplement for "A three-dimensional musculoskeletal model of the dog"

#### Mass

The volume data from Amit et al (2009) and CT-data from Andrada (unpub. data) were used to calculate the mass of the segments. The individual segments were calculated as percentages of the total body weight of the Beagle Simon (13.8 kg).

#### Inertia

For the calculation of the moment of inertia, the function "calculate.inertia.points" of the free program cloud2 was used. This function determines the symmetric inertia matrix for the individual segments on the basis of the point clouds of the bone geometries and the segment masses. A complete segmentation of a calibrated CT data set was not available.

**Table inertia**: Moment of inertia of the humerus segment of a beagle based on A) Andrada et al. 2017 B) Bone geometries C) Segmentation of CT data.

| A) Moment of Inertia about CoM $10^{-3}$ [kg m²] | | |
|---|---|---|
| 0.435 | | |
| | 0.329 | |
| | | 0.254 |

| B) Moment of Inertia about CoM $10^{-3}$ [kg m²] | | |
|---|---|---|
| 0.513 | 0.005 | -0.003 |
| | 0.500 | 0.031 |
| | | 0.044 |

| C) Moment of Inertia about CoM $10^{-3}$ [kg m²] | | |
|---|---|---|
| 0.332 | 0.001 | 0.001 |
| | 0.256 | 0.035 |
| | | 0.178 |

The influence of asymmetrical mass distribution was tested using the humerus. The humerus with the surrounding musculature was segmented and the moment of inertia was calculated with the function "scalar.inertia" of the free software imagexd. The comparison showed that the component Izz of the calculation from the bone geometries showed the greatest difference (Tab. inertia). This can be explained by the fact that the real geometries along the X and Y axis were neglected. However, the main moment is determined by the longitudinal direction (bone length). The change between tensors in simulations did not significantly influence either joint-torque nor muscle activation profiles.

#### Inverse dynamics (3D-Newton Euler Method)

This method is based on Winter 2010, and has been presented in Andrada et al., 2017. Synchronized 3-D kinematics, kinetics, and segmental properties were processed and combined in a customized Matlab program. The 3-D coordinates of marker trajectories were smoothed using a fourth-order Butterworth low-pass filter with a cutoff frequency of 6 Hz. To obtain 3-D angular kinematics, a Cardan sequence of 3 rotations around the $x$, $y$, and $z$ axes was used as described. The transformation matrix for the Cardan sequence was as follows:

$$\begin{bmatrix} x_3 \\ y_3 \\ z_3 \end{bmatrix} = \begin{bmatrix} c_2c_3 & s_3c_1 + s_1s_2c_3 & s_1s_3 - c_1s_2c_3 \\ -c_2c_3 & c_1c_3 - s_1s_2s_3 & s_1c_3 + c_1s_2s_3 \\ s_2 & -s_1c_2 & c_1c_2 \end{bmatrix} \begin{bmatrix} x_0 \\ y_0 \\ z_0 \end{bmatrix}$$

where the indices 1,2,3 indicate the angles $\theta_1$, $\theta_2$, $\theta_3$. For example c1 means cos ($\theta_1$). The angles $\theta_1$, $\theta_2$, $\theta_3$ can thus be obtained from the trigonometric relations present in the transformation matrix. Those angles and their time derivatives are expressed in global coordinates. For the purpose of this study, we were interested in segmental (body-frame) kinematics and dynamics. To transform velocities expressed in laboratory frame into body frame, the following transformation matrix was used:

$$\begin{bmatrix} \omega_x \\ \omega_y \\ \omega_z \end{bmatrix} = \begin{bmatrix} c_2c_3 & s_3 & 0 \\ -c_2c_3 & c_3 & 0 \\ s_2 & 0 & 1 \end{bmatrix} \begin{bmatrix} \dot{\theta}_1 \\ \dot{\theta}_2 \\ \dot{\theta}_3 \end{bmatrix}$$

where the angular accelerations in body frame ($\alpha_x$, $\alpha_y$, and $\alpha_z$) are simply the time derivatives of the respective velocities ($\omega_x$, $\omega_y$, and $\omega_z$). Then, the force (resp. torque) around the proximal end of each segment was estimated by use of the Newton-Euler equations of motion in the CoM as follows:

$$R_{xp} = R_{xd} + m\ddot{x}$$

$$R_{yp} = R_{yd} + m\ddot{y}$$

$$R_{zp} = R_{zd} + m\ddot{z} + mg$$

$$M_{xp} = I_1\alpha_x + (I_3 - I_2)\omega_z\omega_y + R_{yd}l_d + R_{yp}l_p + M_{xd}$$

$$M_{yp} = I_2\alpha_y + (I_1 - I_3)\omega_x\omega_z + R_{xd}l_d + R_{xp}l_p + M_{yd}$$

$$M_{zp} = I_3\alpha_z + (I_2 - I_1)\omega_y\omega_x + M_{zd}$$

where $m$ is the mass of the segment, $I$ is the principal moment of inertia of the segment around the CoM, $R$ is the reaction force, $M$ is the torque in the proximal (p) and distal (d) ends of the segment, and $l$ is the lever arm of the forces relative to the CoM.

#### Tables

**Table S1:** Complete list of all muscles for the whole model and for the sub-models (fore- and hindlimb ⇨ X). The number of sub muscles is shown in brackets.

| | **forelimb** | **hindlimb** |
|---|---|---|
| M. trapezius cervical | X | – |
| M. trapezius thoracic | X | – |
| M. rhomboideus thoracic | X | – |
| M. rhomboideus cervicis | X | – |
| M. latissimus dorsi | X | – |
| M. brachiocephalicus [1 - 2] | X | – |
| M. serratus [1 - 12] | X | – |
| M. pectoralis profundus | X | – |
| M. pectoralis superficialis | X | – |
| M. supraspinatus | X | – |
| M. infraspinatus | X | – |
| M. subscapularis | X | – |
| M. deltoideus sca | X | – |
| M. deltoideus acr | X | – |
| M. teres major | X | – |
| M. teres minor | X | – |
| M. coracobrachialis | X | – |
| M. biceps brachii | X | – |
| M. brachialis | X | – |
| M. triceps caput longum | X | – |
| M. triceps caput laterale | X | – |
| M. triceps caput accessorium | X | – |
| M. triceps caput medial | X | – |
| M. anconeus | X | – |
| M. pronator teres | X | – |
| M. supinator | X | – |
| M. flexor carpi radialis | X | – |
| M. flexor carpi ulnaris | X | – |
| M. extensor carpi ulnaris | X | – |

| | | |
|---|---|---|
| M. extensor carpi radialis | X | – |
| M. abductor policis longum | X | – |
| M. longissimus | – | – |
| M. quadratus lumborum | – | – |
| M. iliocostalis | – | – |
| M. sacrocaudalis | – | – |
| M. psoas major | – | X |
| M. psoas minor | – | X |
| M. adductor longus | – | X |
| M. gemelli | – | X |
| M. gluteus medius | – | X |
| M. gluteus profundus | – | X |
| M. gluteus superficialis | – | X |
| M. iliacus | – | X |
| M. obturator ext | – | X |
| M. obturator int | – | X |
| M. piriformis | – | X |
| M. pectineus | – | X |
| M. sartorius caudal | – | X |
| M. sartorius cranial | – | X |
| M. tensor fasciae | – | X |
| M. quadratus femoris | – | X |
| M. vastus lateralis | – | X |
| M. vastus medialis | – | X |
| M. rectus femoris | – | X |
| M. abductor cruris | – | X |
| M. adductoren | – | X |
| M. biceps femoris longum | – | X |
| M. biceps femoris breve | – | X |
| M. gracilis tractus calcaneus | – | X |
| M. semimembranosus | – | X |
| M. semitendinosus | – | X |
| M. gastrocnemius lateralis | – | X |
| M. gastrocnemius medialis | – | X |

| | | |
|---|---|---|
| M. peroneus longus | | X |
| M. tibialis caudalis | | X |
| M. tibialis cranialis | | X |
| M. popliteus | | X |

### Figures

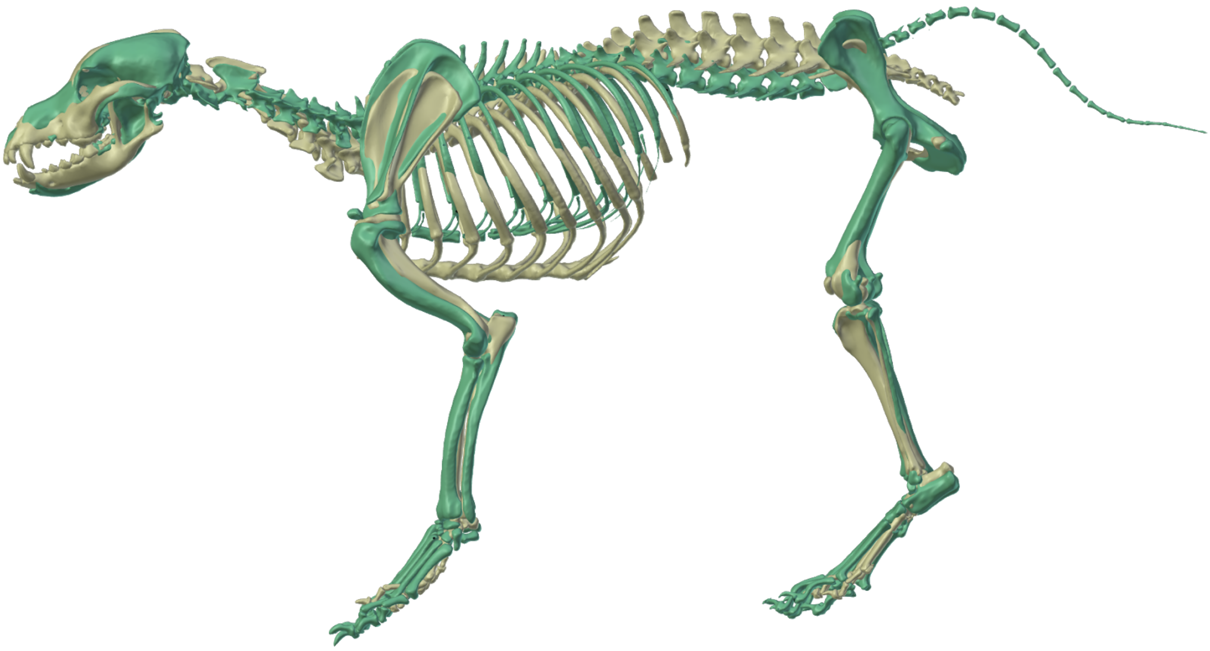

**Figure S1:** Visualization of the reconstructed bones of the GS-model (grey) and the BE-model (green). The BE model was scaled to the size of the GS model.

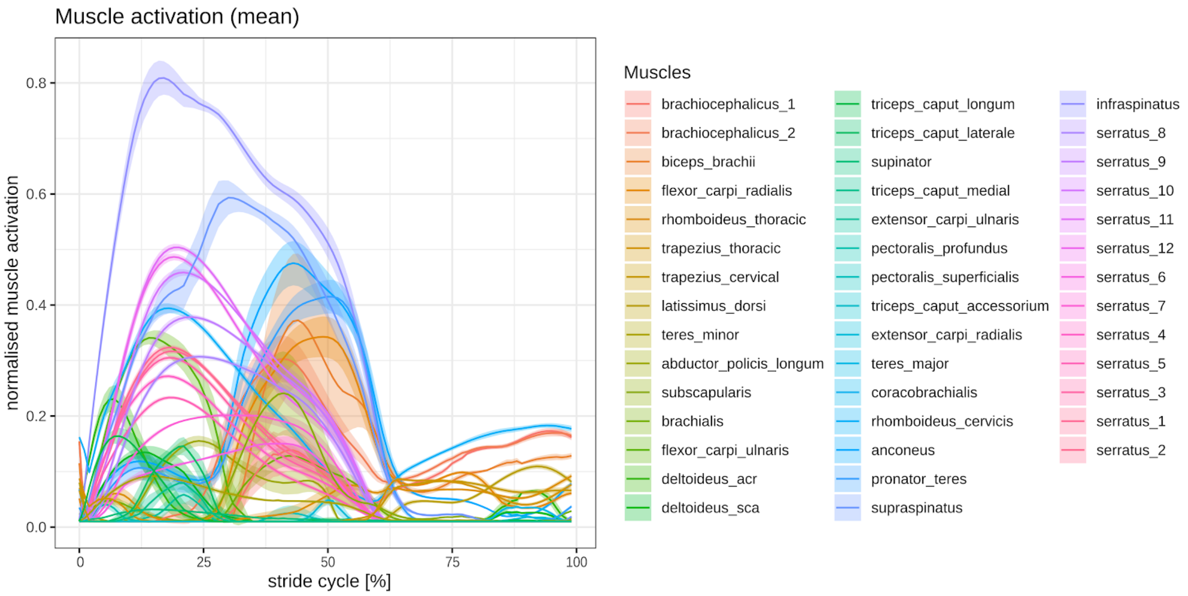


**Figure S2:** Simulated forelimb muscle activation in a walking beagle for 43 muscles or muscle parts. The standard deviation is shown as shaded bands.

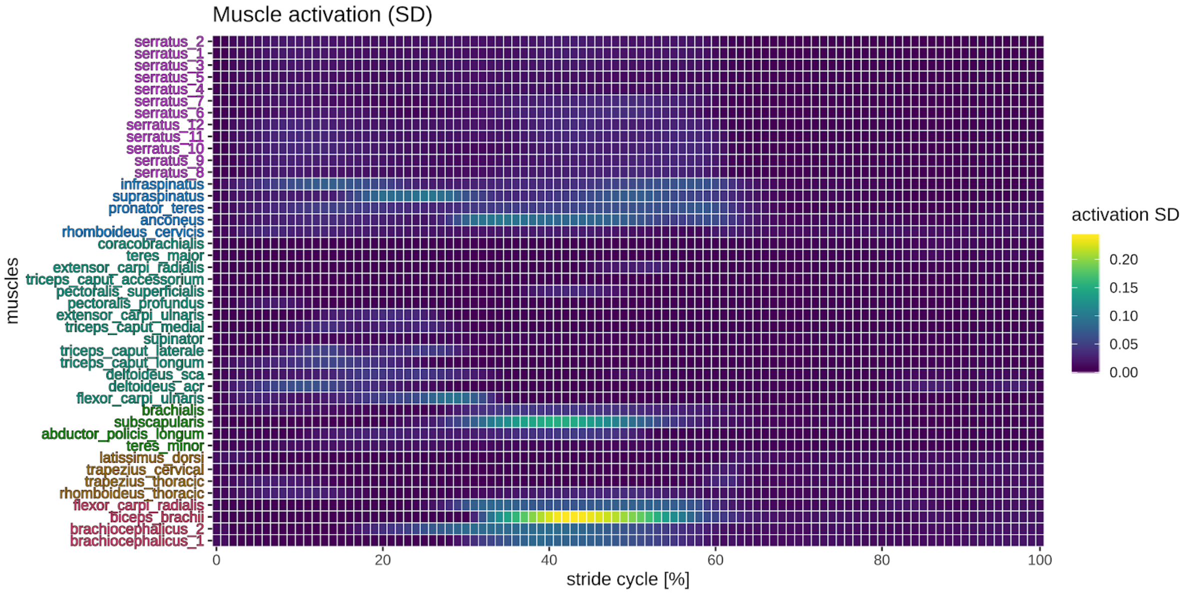


**Figure S3:** Simulated forelimb muscle activation in a walking beagle. Logarithmic muscle SD is shown from low (purple) to high (yellow).

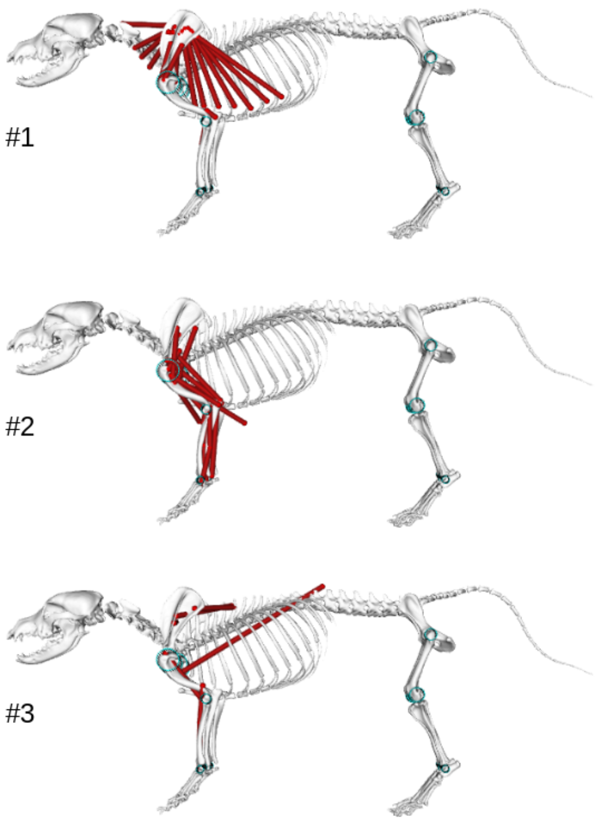


**Figure S4:** Representation of the muscles according to hierarchical clustering (method – ward.d2) and minimal leaf sorting of simulated forelimb logarithmic muscle activation in the walking beagle.

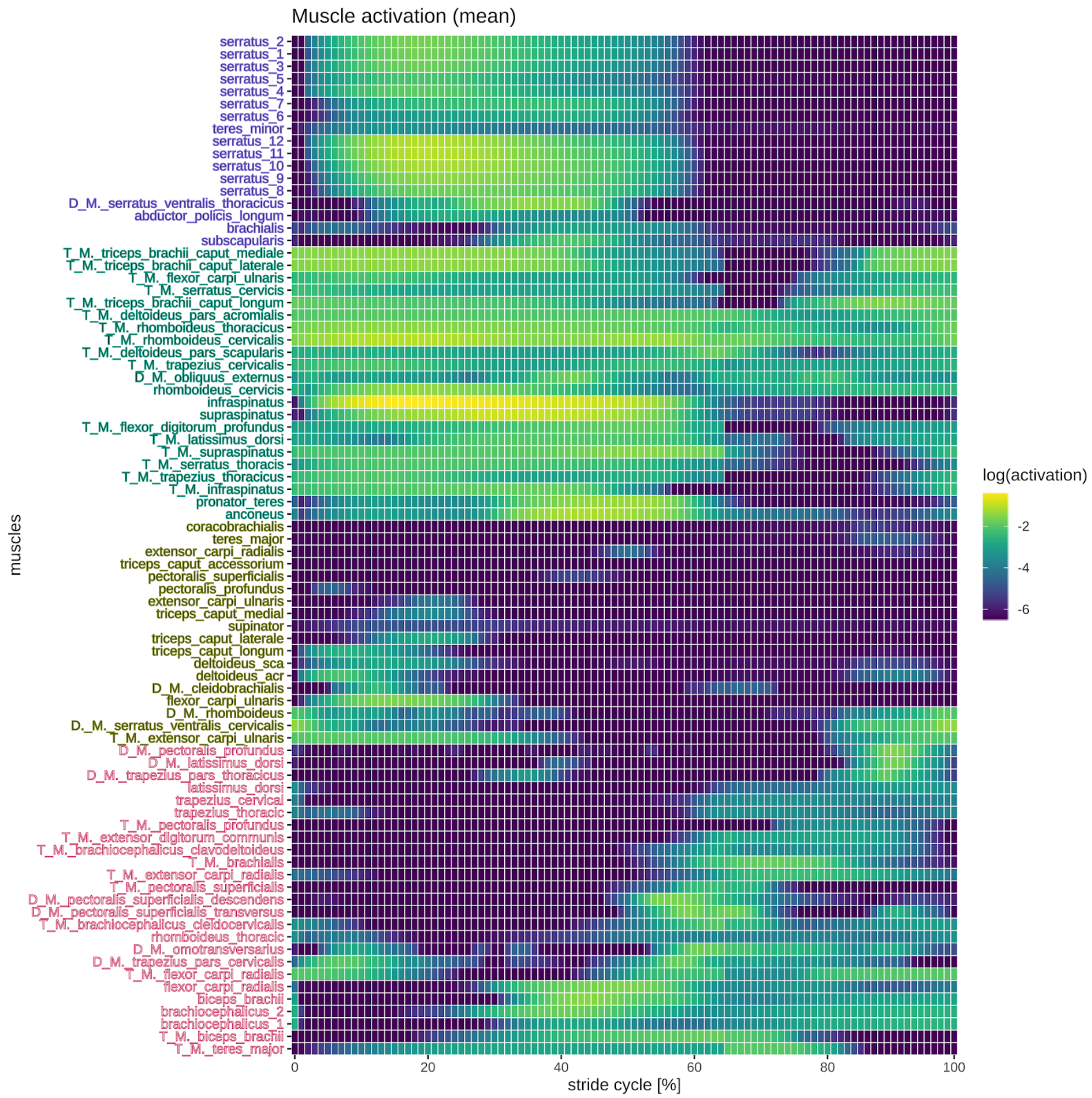


**Figure S5:** Simulated and literature (T: Tokuriki, 1973; D: Deban et al., 2012) forelimb muscle activation in a walking beagle. Logarithmic muscle activation is shown from low (purple) to high (yellow).

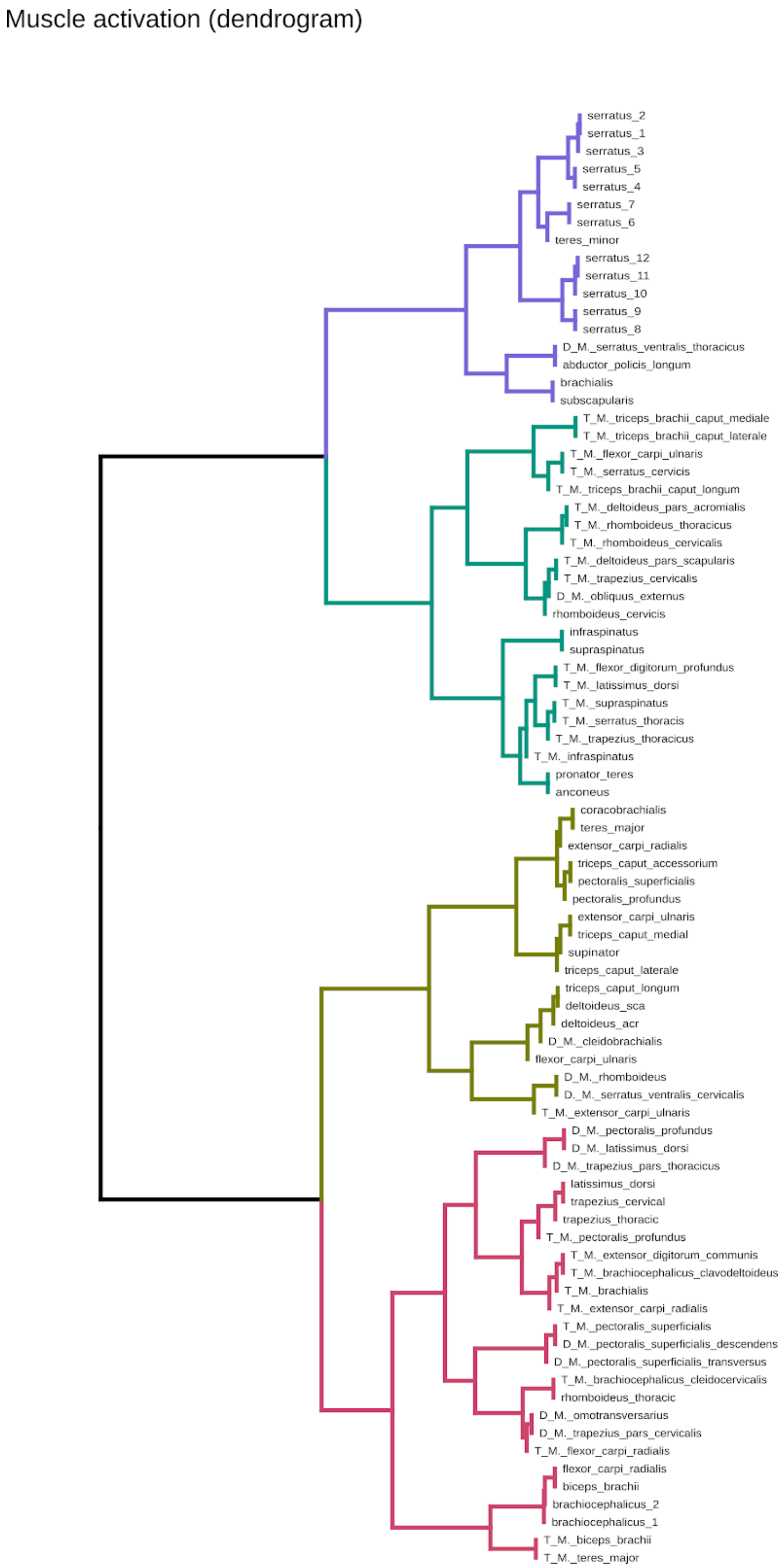


**Figure S6:** Hierarchical clustering (method – ward.d2) and minimal leaf sorting of simulated and literature (T: Tokuriki, 1973; D: Deban et al., 2012) forelimb logarithmic muscle activation in a walking beagle.
